# pH Gradient Mitigation in the Leaf Cell Secretory Pathway Alters the Defense Response of *Nicotiana benthamiana* to Agroinfiltration

**DOI:** 10.1101/431767

**Authors:** Philippe V. Jutras, Frank Sainsbury, Marie-Claire Goulet, Pierre-Olivier Lavoie, Rachel Tardif, Louis-Philippe Hamel, Marc-André D’Aoust, Dominique Michaud

## Abstract

Partial neutralization of the Golgi lumen pH by ectopic expression of influenza virus M2 proton channel stabilizes acid-labile and protease-susceptible recombinant proteins in the plant cell secretory pathway. Here, we assessed the impact of M2 channel expression on the proteome of *Nicotiana benthamiana* leaf tissue infiltrated with the bacterial gene vector *Agrobacterium tumefaciens*, keeping in mind the key role of pH homeostasis on secreted protein processing and the involvement of protein secretion processes in plant cells upon microbial challenge. The proteomes of leaves agroinfiltrated with an empty vector or with an M2 channel-encoding vector were compared with the proteome of non-infiltrated leaves using a iTRAQ quantitative proteomics procedure. Leaves infiltrated with the empty vector had a low soluble protein content compared to non-infiltrated leaves, associated with a strong decrease of photosynthesis-associated proteins (including Rubisco) and a parallel increase of stress-related secreted proteins (including pathogenesis-related proteins, protease inhibitors and molecular chaperones). M2 expression partly compromised these alterations of the proteome to restore original soluble protein and Rubisco contents, associated with higher levels of translation-associated (ribosomal) proteins and reduced levels of stress-related proteins in the apoplast. Proteome changes in M2-expressing leaves were determined both transcriptionally and post-transcriptionally, to alter the steady-state levels of proteins not only along the secretory pathway but also in other cellular compartments including the chloroplast, the cytoplasm, the nucleus and the mitochondrion. These data illustrate the cell-wide influence of Golgi lumen pH homeostasis on the leaf proteome of *N. benthamiana* plants responding to microbial challenge. They underline in practice the relevance of carefully considering the eventual off-target effects of accessory proteins used to modulate specific cellular or metabolic functions in plant protein biofactories.

## INTRODUCTION

Major advances in plant cell biology and genetic engineering have bolstered the use of plants as expression hosts for clinically and industrially valuable recombinant proteins (Stöger et al., 2014; Sack et al., 2015; Lomonossoff and D’Aoust, 2016; Tschofen et al., 2016). An array of DNA vectors, regulatory sequences and delivery systems have been developed for high-level transgene expression in plant systems (Streatfield, 2007; Makhzoum et al., 2014). Basic knowledge on protein biosynthetic pathways in plants has been translated in parallel to plant expression hosts, helpful to sustain proper maturation of expressed proteins or to implement novel cellular functions for effective protein processing *in planta* (Faye et al., 2005; Gomord et al., 2010; Mandal et al., 2016). Current efforts to further strengthen the position of plants as valuable protein expression hosts also include the development of metabolic, cellular or phenology engineering approaches to address specific issues related to recombinant protein maturation, stability or recovery. Examples are the ectopic implementation of a dwarf plant phenotype to optimize culture area use in the greenhouse (Nagatoshi et al., 2015), activation of the octadecanoid pathway to reduce ribulose 1,5-*bis*-phosphate carboxylase oxygenase (Rubisco) loads in leaves prior to protein purification (Robert et al., 2015), or the expression of accessory convertases to generate biologically active forms of therapeutic proteins otherwise requiring chemical refolding after extraction (Wilbers et al., 2016). Other examples are the expression of protease inhibitors to prevent unintended protein degradation by resident proteases (Goulet et al., 2012; Pillay et al., 2014; Jutras et al., 2016), or the expression of an accessory proton channel to stabilize acid-labile and proteolysis-susceptible proteins in the Golgi lumen (Jutras et al., 2015; 2018).

A key challenge now to harness the full potential of these emerging engineering approaches and to take advantage of their eventual synergistic effects *in planta* is to decipher their possible ‘off-target’ effects *in planta.* By definition, rational schemes for plant metabolic engineering target specific physiological or enzymatic processes but unintended effects in the modified host cannot be ruled out, especially in those cases where the ectopic effectors alter basic cellular functions or physicochemical parameters. For instance, the inhibition of host endogenous proteases with accessory protease inhibitors is useful to prevent the degradation of protease-susceptible recombinant proteins *in planta* (Robert et al., 2016) but interfering effects on protein biosynthetic and turnover rates in the modified host could in some cases have an impact, positive or negative, on overall protein yields recovered from source tissues (Badri et al., 2009; Goulet et al., 2010a). Similarly, partial neutralization of the Golgi lumen pH by ectopic expression of influenza virus M2 proton channel (Holsinger et al., 1994) is useful to stabilize acid-labile proteins *in situ* (Jutras et al., 2015) but eventual effects of this transporter on host physiological functions remain likely given the influence of pH homeostasis on protein posttranslational maturation, processing and trafficking along the cell secretory pathway (Schumacher, 2014; Jutras et al., 2018).

Our goal in this study was to assess the impact of M2 channel expression on the leaf proteome of *Nicotiana benthamiana* infiltrated with the bacterial gene vector *Agrobacterium tumefaciens.* M2 forms tetrameric transmembrane channels for proton extrusion in the cytosol of infected mammalian cells, to generate an increased pH in the Golgi lumen favourable to the folding and stability of the influenza virus glycoproteins (Schnell and Chou, 2008; Cady et al., 2009). Transient expression of M2 in *N. benthamiana* leaves was shown to trigger a similar pH increase in the *cis*-and trans-Golgi compartments, useful to stabilize acid-labile recombinant proteins and peptide linkers migrating towards to the apoplast (Jutras et al., 2015). A side effect of the viral transporter was also reported recently, by which the activity of pH-dependent resident proteases and their impact on the integrity of protease-susceptible proteins in the secretory pathway is altered upon pH increase (Jutras et al., 2018). An unsolved question at this point is to what extent M2 expression exerts pleiotropic effects on host plant cellular functions via its primary effect on pH homeostasis in the secretory pathway. As in other eukaryotic cells (Orlowski and Grinstein, 2011), a pH gradient is naturally established in plant cells between the ER and the Golgi (Martinière et al., 2013; Shen et al., 2013) by the combined action of V-type H^+^-ATPases for lumen acidification and Na^+^/H^+^ antiporters for proton efflux and pH fine-tuning (Bassil and Blumwald, 2014). Early transfection studies with animal cell models showed significant effects of M2 ectopic expression on endogenous protein trafficking and Golgi cisternae morphology, presumably due to a disturbed balance of ‘in and out’ proton movements through the Golgi membranes (Sakaguchi et al., 1996; Henkel et al., 1998). Similarly, altered pH in the Golgi of Arabidopsis Na^+^/H^+^ antiporter knockouts, or in the Golgi of plant cells treated with chemical inhibitors of V-type H^+^-ATPases or Na^+^/H^+^ antiporters, was associated with altered rates of protein posttranslational processing and trafficking towards the late secretory pathway compartments (Dettmer et al., 2006; Martinière et al., 2013; Ashnest et al., 2015; Reguera et al., 2015; Wu et al., 2016). We here followed an ‘isobaric tags for relative and absolute quantification’ (iTRAQ) liquid chromatography (LC)-mass spectrometry (MS/MS) approach (Brewis and Brennan, 2010) to probe the influence of Golgi lumen pH homeostasis on the whole proteome of agroinfiltrated *N. benthamiana* leaves upon M2 expression, keeping in mind the significance of protein secretion and trafficking in plant cells responding to microbial challenge (Inada and Ueda, 2014; Ben Khaled et al., 2015).

## RESULTS AND DISCUSSION

### Agroinfiltration Alters the Leaf Proteome of *N. benthamiana*

Total soluble proteins and total numbers of up- and downregulated proteins in whole-cell samples were first determined to get a general overview of proteome alterations in leaves infiltrated either with agrobacteria harboring an M2 channel-encoding vector, or with agrobacteria harboring an ‘empty’ version of the same vector (EV), to measure the effects of bacterial infiltration and M2 expression on leaf protein content at the cell-wide scale (**Fig. 1**). A significant adjustment of the protein complement was suggested by a low total soluble protein content of 5.7 mg.g^−1^ fresh weight in EV-infiltrated leaves compared to 9.4 mg.g^−1^ in non-infiltrated leaves 6 d post-infiltration (post-ANOVA LSD; *P*<0.05) (**Fig. 1A**). Protein content reduction in agroinfiltrated leaves was associated with a downregulation of ribulose-1,5-*bis*-phosphate carboxylase/oxygenase (Rubisco) large and small subunits and a concomitant upregulation of protein bands in the 20–35-kDa range (**Fig. 1B**). By contrast, protein content in M2 vector-infiltrated leaves was estimated at 8.4 mg.g^−1^ fresh weight, statistically similar to non-infiltrated leaves (LSD; *P*>0.05) (**Fig. 1A**). Rubisco subunit band intensities were also comparable in M2 vector-infiltrated and non-infiltrated leaves, two to three times more intense than the corresponding bands in EV-infiltrated leaves (**Fig. 1B**). A time-course analysis over 12 d post-infiltration indicated a rapid and durable upregulating effect of M2 expression on soluble protein content compared to EV-infiltrated leaves, already measurable after 2 d and still important after 12 d (**Supplemental Fig. S1**). No positive effect on protein content was observed in leaves infiltrated to express ^A30P^M2, an inactive single mutant of M2 (Holsinger et al., 1994) (**Supplemental Fig. S1**), indicating a link between the protein content-restoring effect of the transporter and its proton-conducting activity in agroinfiltrated leaves.

**Figure 1.**
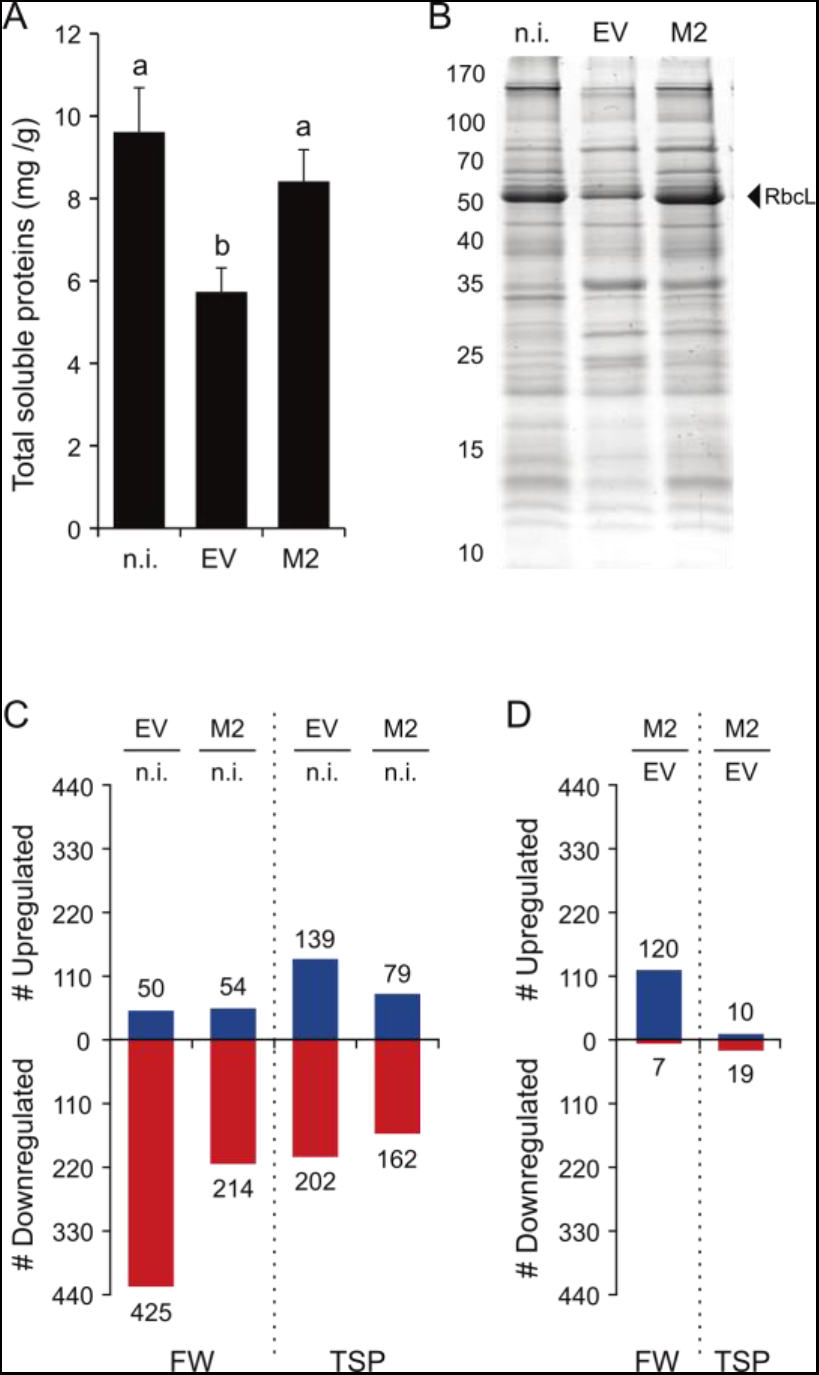
Soluble protein content and proteome changes in non-infiltrated (n.i.), EV-infiltrated and M2 channel-expressing leaves 6 d post-agroinfiltration. (A) Total soluble proteins (TSP) in leaf tissue, as expressed on a leaf fresh weight basis. Data are the mean of three biological (plant) replicate values ± sd. Bars with the same letter are not significantly different (post-ANOVA LSD; *P*<0.05). (B) Soluble protein profiles in leaves as observed on Coomassie blue-stained polyacrylamide gels following 12% w/v SDS-PAGE. Numbers on the left refer to commercial molecular weight markers. Arrow on the right points to the large, 52-kDa subunit of Rubisco. (C) Numbers of iTRAQ-identified proteins up- (blue) or down- (red) regulated by at least twofold in EV-infiltrated and M2-expressing leaves compared to non-infiltrated control leaves. (D) Numbers of iTRAQ-identified proteins up (blue) or down- (red) regulated (blue) by at least twofold in M2-expressing leaves compared to EV-infiltrated leaves. Data on panels C and D are expressed on a leaf fresh weight (FW) or protein-specific (TSP) basis. Additional data on leaf soluble protein content are provided in **Supplemental Fig. S1.**

We conducted a iTRAQ analysis of non-infiltrated and infiltrated leaf protein extracts to estimate the overall impacts of agroinfiltration and M2 expression on the leaf soluble protein complement (**Fig. 1C,D**) and to characterize the specific effects of these treatments at the cell-wide scale (**Figs. 2–5**). A total of 5,928 unique peptides were detected by MS/MS, allowing for the identification of 2,388 proteins at a confidence level of 95% (*P*<0,05). A little more than 50% (i.e. 1,255) of these proteins were identified based on at least two unique peptides and used for further comparative assessments (**Supplemental Table S1**). On a leaf fresh weight (FW) basis, 425 proteins were downregulated, and 50 proteins upregulated, by at least twofold in EV-infiltrated leaves compared to 214 proteins downregulated and 54 upregulated in M2 vector-infiltrated leaves (**Fig. 1C**). On a total soluble protein (TSP) basis, 202 proteins were downregulated, and 139 upregulated, in EV-infiltrated leaves compared to non-infiltrated leaves, roughly similar to protein numbers flagged as down- (162) or upregulated (79) upon M2 expression (**Fig. 1C**). A pairwise comparison of MS/MS data produced for EV- and M2 vector-infiltrated leaves was performed to estimate the overall impact of M2 proton channel activity on agroinfiltration-induced proteome changes (**Fig. 1D**). On a TSP basis, only 10 proteins were upregulated, and 19 downregulated, by at least twofold in M2 vector-infiltrated leaves compared to EV-infiltrated leaves, out of 1,255 proteins monitored. These data suggested overall a qualitative impact of M2 expression on the host proteome limited, on protein-specific basis, to a relatively small number of up- or downregulated proteins. They confirmed, by contrast, the strong impact of agroinfiltration on the leaf proteome, attenuated to some extent by Golgi pH alteration in M2-expressing cells.

**Figure 2.**
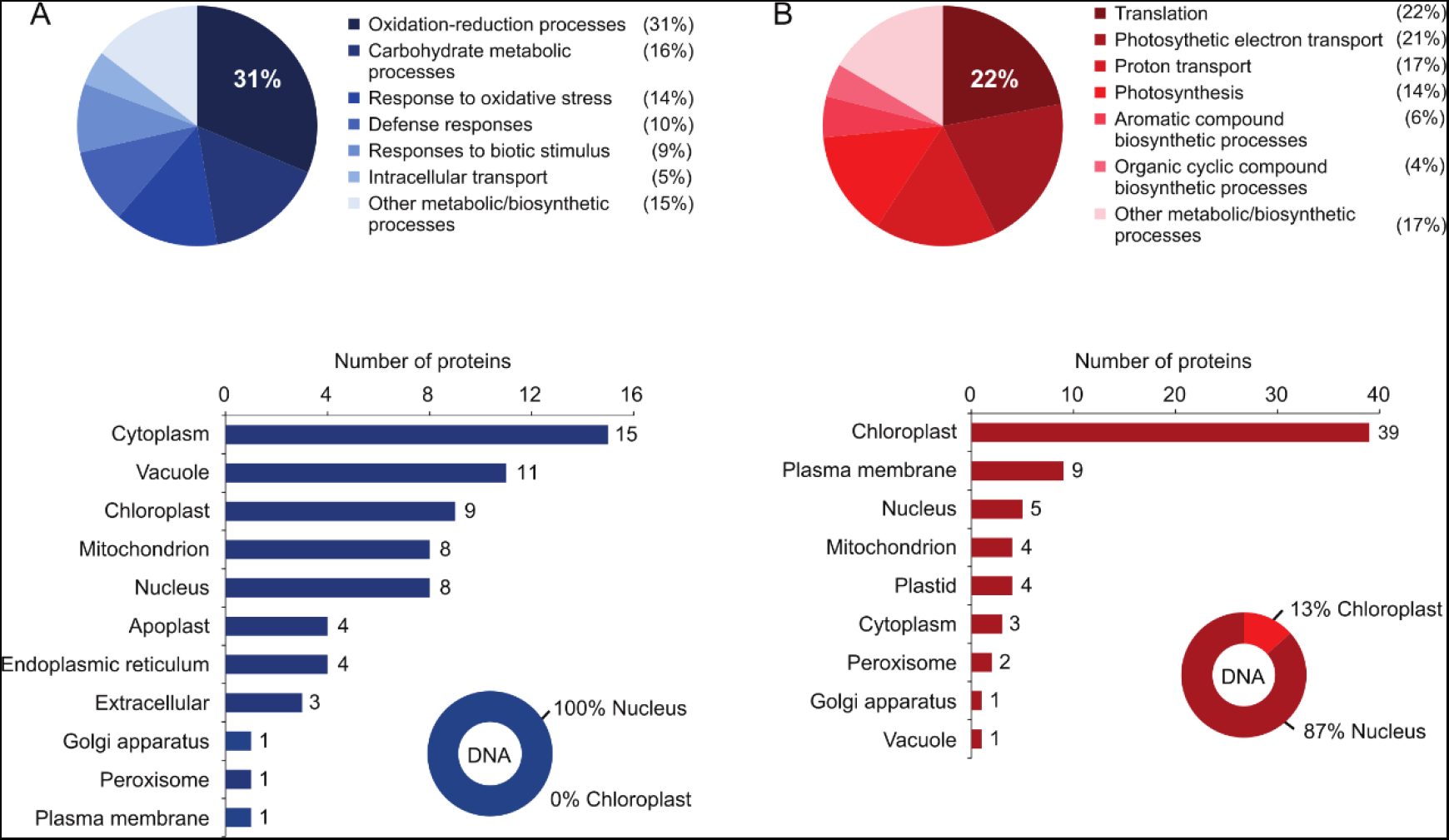
GO enrichment analysis of iTRAQ-quantified proteins up- (A, in blue) or down- (B, in red) regulated by at least twofold in EV-infiltrated leaves compared to non-infiltrated (n.i.) leaves. Pie charts identify the six most affected biological processes in leaves as inferred from biological functions assigned to the 60 most upregulated, or 60 most downregulated, proteins in EV-infiltrated leaves. Bar charts summarize the subcellular distribution of these proteins as inferred *in silico* from their predicted cellular localization. Circle charts indicate the relative abundance of chloroplast genome- and nuclear genome-encoded proteins among these same proteins. The 60 most upregulated, and 60 most downregulated, proteins in EV-infiltrated leaves compared to non-infiltrated leaves are listed in **Supplemental Tables S2** and **S3**, respectively. Complementary information to this GO enrichment analysis is provided in **Supplemental Fig. S2**.

### Agroinfiltration Triggers a Classical Biotrophic Pathogen-Inducible Defense Response in Leaves

We performed BLAST alignments and a Gene Ontology (GO) enrichment analysis of our MS/MS dataset to classify the most significant proteome alterations in agroinfiltrated leaves based on the biological roles, biochemical functions and/or subcellular locations assigned *in silico* to the regulated proteins (**Fig. 2**, **Supplemental Fig. S2**). Agroinfiltration was shown previously to trigger the secretion of stress-related proteins in *N. benthamiana* leaves including salicylic acid-inducible pathogenesis-related (PR) protein PR-1a and several PR-2 (ß-glucanase), PR-3 (chitinase) and PR-5 (osmotin) protein isoforms (Pruss et al., 2008; Goulet et al., 2010b; Robert et al., 2015). We here assigned biological roles and cellular locations to the 60 most upregulated, and 60 most downregulated, proteins in EV-infiltrated plants representing, together, 10% of the proteins confidently identified by MS/MS and 25% of the proteins up- or downregulated by at least twofold in agroinfiltrated leaf tissue. In line with previous studies, a range of defense and oxidative stress-related proteins were induced in EV-infiltrated leaves, including oxidoreductases (e.g. peroxidases), molecular chaperones (e.g. heat shock proteins), PR proteins (e.g. ß-glucanases, chitinases, osmotins) and protease inhibitors (e.g. Kunitz proteins) (**Supplemental Table S2**). Stress-related proteins accounted for approximately two thirds of the upregulated proteins (**Fig. 2A**, upper panel), distributed in several cellular compartments including the chloroplast, the mitochondrion, the nucleus, the cytosol and different parts of the cell secretory pathway (**Fig. 2A**, lower panel). Similar to conclusions drawn earlier for the leaf apoplast proteome (Goulet et al., 2010b), all upregulated proteins in soluble protein samples, including chloroplastic proteins, were nuclear genome-encoded (**Fig. 2A**, lower panel).

Unlike stress-related proteins, several proteins involved in mRNA translation, photosynthesis and ATP biosynthesis were strongly downregulated in EV-infiltrated leaves (**Fig. 2B**, upper panel). Chloroplastic proteins including Rubisco (also see **Fig. 1B**), structural subunits of photosystems I and II, chlorophyll-binding proteins, ATP synthase subunits and translation-associated ribosomal proteins (**Supplemental Table S3**) accounted for more than 75% of the downregulated proteins (**Fig. 2B**, lower panel). A small proportion (13%) of these proteins were chloroplast genome-encoded but the vast majority were encoded by the nuclear genome as observed for the upregulated proteins (**Fig. 2B**, lower panel). These observations supported overall those current experimental models suggesting the onset of growth–defense trade-offs in microbe-infected plants, whereby energy production- and photosynthesis-associated genes are downregulated upon microbial challenge (or ectopic application of salicylic acid) to limit the availability of carbon resources to the invading organism or to promote defense responses over primary metabolism-related processes (Sugano et al., 2010; Takatsuji, 2017). Our data showing an impact of agroinfiltration at the whole cell scale also reminded the strong influence of pathogen infection on nuclear gene expression and the likely implication of leaf chloroplasts as stress signal receivers and pro-defense secondary signal transmitters to the nucleus upon microbial attack (Serrano et al., 2016). Primary signals transmitted into plant cells to induce immune responses following membrane receptor-mediated recognition of pathogen-associated molecular patterns (Boller and Felix, 2009) are readily relayed to the chloroplasts, where they trigger the production of retrograde secondary signals that move towards the nucleus to activate defense-related genes and repress chloroplast protein-encoding genes (Nomura et al., 2012). Retrograde signals identified in recent years were shown to induce nuclear genes involved in the biosynthesis of salicylic acid (Nomura et al., 2012; Xiao et al., 2012; Ishiga et al., 2017), a key elicitor of immune responses to agroinfiltration in *N. benthamiana* leaves (Anand et al., 2008; Pruss et al., 2008).

### M2 Channel Expression Attenuates the Host Plant Response to Agroinfiltration

A complementary GO enrichment analysis was performed to characterize eventual interfering effects of M2 expression on host leaf proteome adjustments upon agroinfiltration (**Fig. 3**, **Supplemental Fig. S3**). Our observations above about the overall effects of EV and M2 vector infiltrations (**Fig. 1**) suggested a limited impact of M2 on the proteome of agroinfiltrated plants as expressed in numbers of proteins up- or downregulated by at least twofold in leaf tissue. By contrast, they indicated a strong positive effect of the viral transporter on soluble leaf protein and Rubisco contents that suggested a possibly attenuated defense response upon infection associated with an alteration of pH gradient homeostasis along the cell secretory pathway. We here addressed these questions by comparing the proteomes of EV- and M2 vector-infiltrated leaf protein samples, considering the 60 most upregulated, and 60 most downregulated, proteins in M2-expressing leaves relative to EV-infiltrated leaves taken as a control. Supporting the hypothesis of a defense-attenuating effect for M2, chloroplast and nuclear genome-encoded proteins downregulated in EV-infiltrated leaves were found at higher levels in M2-expressing leaves (**Fig. 3A**) including, along with Rubisco (**Fig. 1**), a range of chloroplastic proteins involved in photosynthesis, ATP biosynthesis and mRNA translation (**Supplemental Table S4**). Likewise, nuclear genome-encoded proteins upregulated in leaves upon EV infiltration including PR proteins, Kunitz protease inhibitors and stress-related oxidoreductases (**Supplemental Table S5**), were found at lower levels in M2-expressing leaves (**Fig. 3B**).

**Figure 3.**
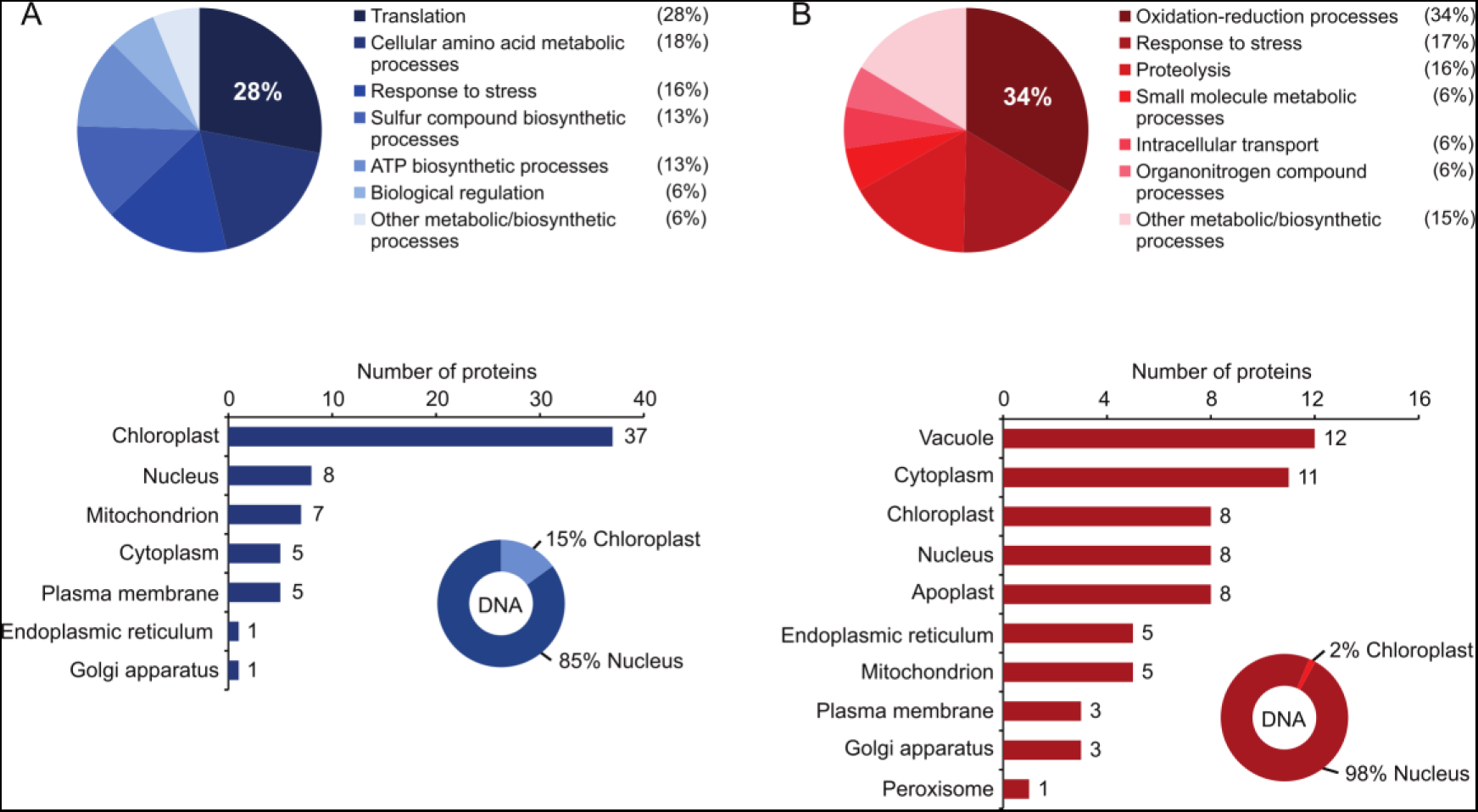
GO enrichment analysis of iTRAQ-quantified proteins up- (A, in blue) or down- (B, in red) regulated by at least twofold in M2 vector-infiltrated leaves compared to EV-infiltrated leaves. Pie charts identify the six most affected biological processes in leaves as inferred from biological functions assigned to the 60 most upregulated, or 60 most downregulated, proteins in M2-expressing leaves. Bar charts summarize the subcellular distribution of these proteins as inferred *in silico* from their predicted cellular localization. Circle charts indicate the relative abundance of chloroplast genome- and nuclear genome-encoded proteins among these same proteins. The 60 most upregulated, and 60 most downregulated, proteins in M2-expressing leaves compared to non-infiltrated leaves are listed in **Supplemental Tables S4** and **S5**, respectively. Additional information on this GO enrichment analysis is provided in **Supplemental Fig. S3**.

Venn diagrams were produced, and a principal component analysis (PCA) performed, to visually compare the proteomes of non-infiltrated and infiltrated leaves (**Fig. 4, Fig. 5**). Of the 60 most upregulated proteins in EV-infiltrated leaves compared to non-infiltrated leaves, 41 (i.e. 68%) were also upregulated in M2-expressing leaves, including several oxidoreductases, PR proteins (ß-glucosidases, chitinases) and ER stress-associated proteins (chaperones, protein disulfide isomerases) (blue circles on **Fig. 4A**). Of the 60 most downregulated proteins in EV-infiltrated leaves, 49 (i.e. 82%) were also downregulated in M2-expressing leaves, including photosynthesis-associated proteins (structural components of photosystem I and II, chlorophyll-binding proteins), protein elongation factors and ATPase complex subunits (red circles on **Fig. 4A**). By comparison, only 18 proteins were found at higher levels, and 29 proteins at lower levels, in both non-infiltrated and M2-expressing leaves compared to EV-infiltrated leaves (**Fig. 4B**). In line with these figures, a PCA analysis of the 1,255 proteins confidently identified by MS/MS in leaf extracts (**Supplemental Table S1**) revealed strongly divergent proteomes in non-infiltrated and EV-infiltrated leaves, compared to M2-expressing leaves exhibiting a hybrid, intermediate proteome (**Fig. 5**). A closer look at the PCA protein distribution indicated a well-defined separation of the 100 most abundant proteins in non-infiltrated and EV-infiltrated leaves (**Supplemental Tables S6 and S7**), unlike the 100 most abundant proteins of EV-infiltrated and M2-expressing leaves (**Supplemental Tables S7** and **S8**) showing a significantly matching distribution (**Fig. 5**). Together, these data confirmed the occurrence of a hybrid, intermediate proteome in M2-expressing leaves, determined first by the host plant defense response to agroinfiltration, and then by an attenuating effect of M2 on this response presumably associated with an alteration of pH homeostasis along the cell secretory pathway.

**Figure 4.**
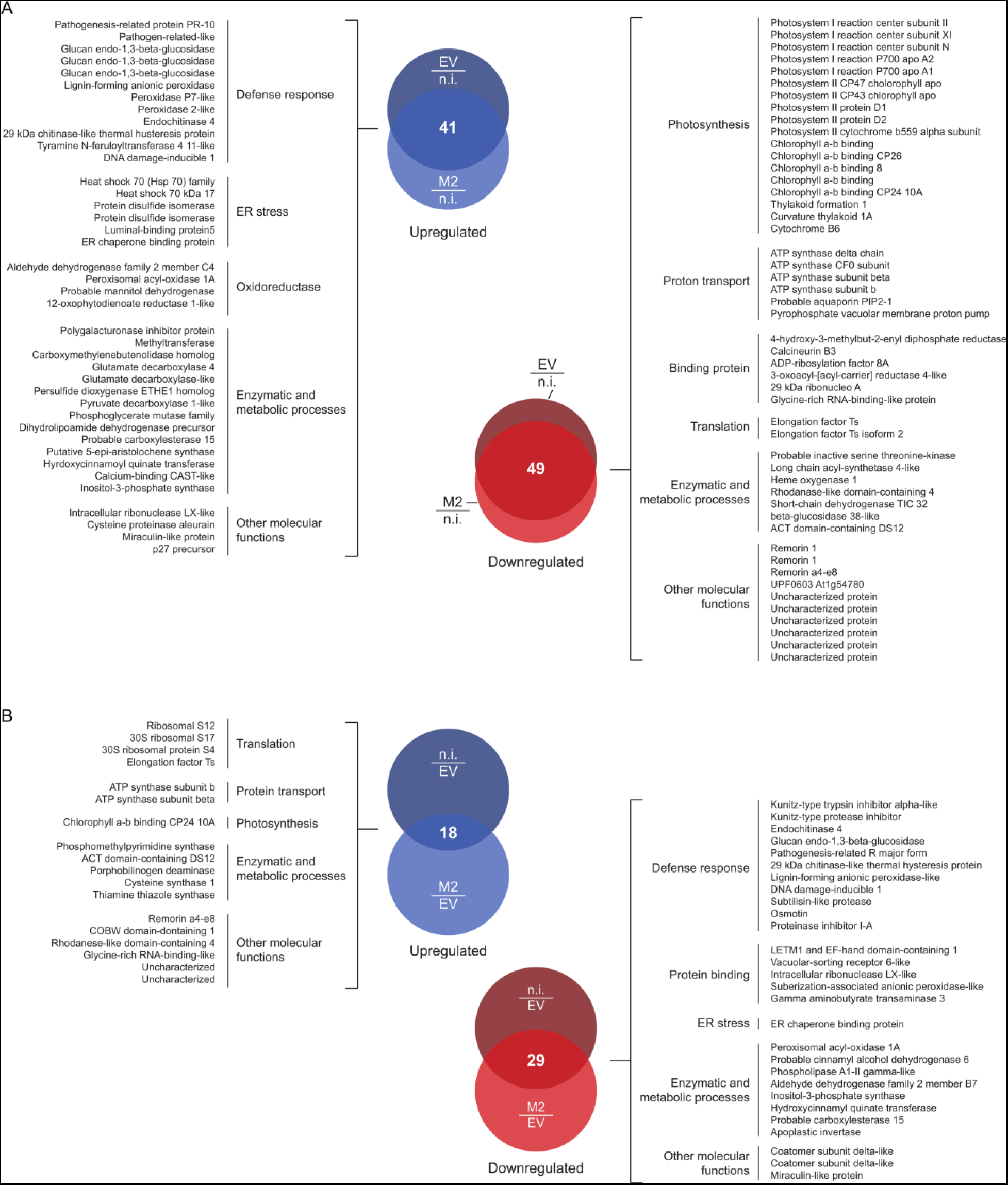
Venn diagrams for the proteins up- (blue) and down- (red) regulated in two [test] treatments relative to the third [reference] treatment. (A) Proteins up- or downregulated in EV-infiltrated and M2-expressing leaves compared to non-infiltrated (n.i.) leaves. (B) Proteins up- or downregulated in non-infiltrated and M2-expressing leaves compared to EV-infiltrated leaves. Numbers in overlapping areas indicate the numbers of proteins up- or downregulated in both test treatments compared to the reference treatment, out of the 60 most affected proteins in each test treatment. Protein lists identify those up- (left-hand side) and down- (right-hand side) regulated proteins shared by the two test treatments.

**Figure 5.**
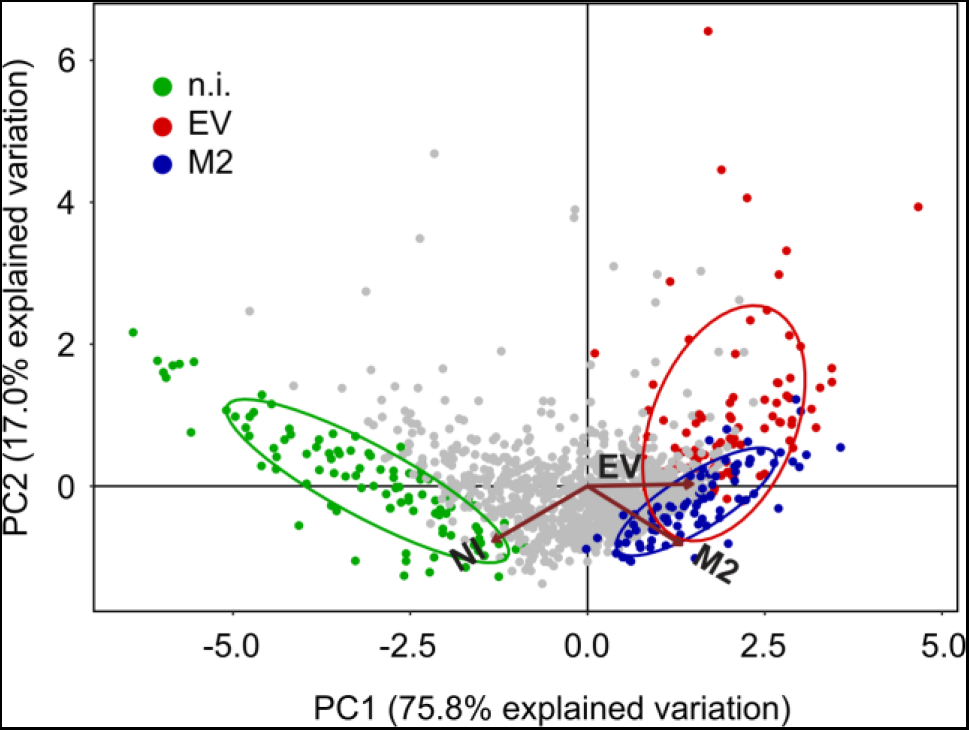
Principal component analysis (PCA) of MS/MS identified proteins in non-infiltrated (n.i.), EV-infiltrated and M2-expressing leaves. The vectors indicate strongly divergent proteomes in non-infiltrated and EV-infiltrated leaves, compared to M2-expressing leaves exhibiting a hybrid proteome matching in part the proteome of EV-infiltrated leaves. The 100 most abundant proteins of each group are coloured to further highlight differences and similarities between non-infiltrated (green), EV-infiltrated (red) and M2-expressing (blue) leaf proteomes. Confidence ellipses show normal data probability for each group of proteins (by default to 68%). The 100 most abundant proteins for either treatments are listed and their relative abundance in leaves given in **Supplemental Tables S6, S7** and **S8**.

### Proteome Alterations in Agroinfiltrated Leaves Are Transcriptionally and Post-Transcriptionally Determined

Immunoblotting and reverse transcriptase (RT)-qPCR analyses were conducted to statistically confirm the validity of our proteomic inferences and to determine whether proteome changes in leaves upon EV infiltration or M2 ectopic expression were transcriptionally or posttranscriptionally regulated (**Fig. 6, Fig. 7**). In line with the Coomassie blue-stained gels above (**Fig. 1**), immunoblot signals for the large and small subunits of Rubisco were the most intense in non-infiltrated leaf samples and the least intense in EV-infiltrated leaf samples (post-ANOVA LSD, *P*<0.05) (**Fig. 6A**). Likewise, PR-2 and PR-3 proteins were readily detected in both EV- and M2 vector-infiltrated leaf samples, unlike non-infiltrated leaves showing no detectable signals on nitrocellulose membranes (*P*<0.05) (**Fig. 6B**). As expected given the well described repressing effects of salicylic acid and microbial challenge on the expression of photosynthesis-associated genes (Shimizu et al., 2007; Sugano et al., 2010), mRNA transcript numbers for the two Rubisco subunits were low in agroinfiltrated leaves compared to non-infiltrated leaves (post-ANOVA LSD, *P*<0.05) (**Fig. 7A**). As also expected, transcript numbers for different stress-related proteins –including PR-3 and PR-10 protein isoforms, ER chaperone-associated protein BIP1 and protein disulfide isomerase PDI7–showed increased levels in agroinfiltrated leaves (*P*<0.05) (**Fig. 7B**).

**Figure 6.**
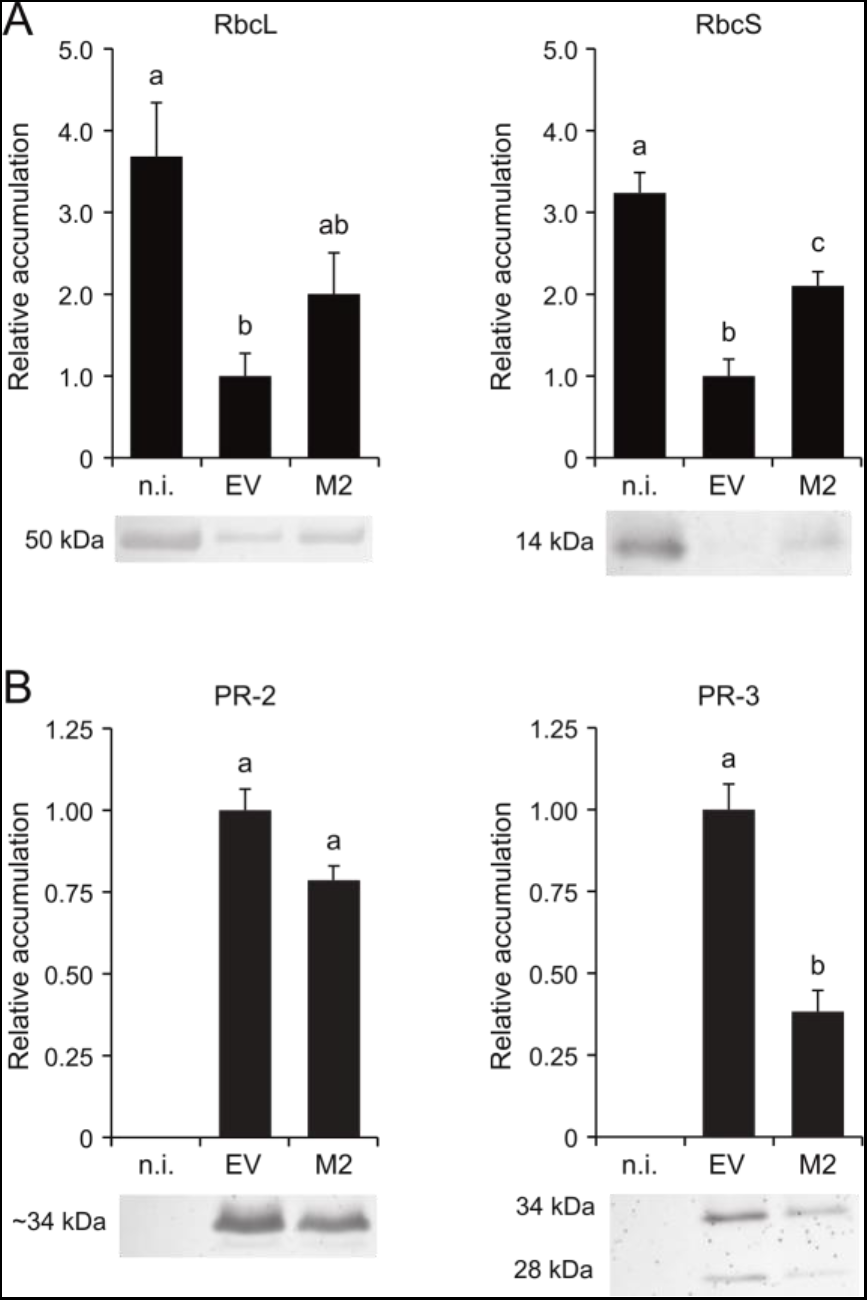
Relative abundance of Rubisco subunits and PR proteins in non-infiltrated (n.i.), EV-infiltrated and M2-expressing leaves. (A) Relative abundance of Rubisco large (RbcL) and small (RbcS) subunits. (B) Relative abundance of PR-2 (ß-glucanase) and PR-3 (chitinase) isoforms. Data are expressed relative to EV-infiltrated leaves (arbitrary value of 1). Each bar is the mean of three biological (plant) replicate values ± se. Bars with the same letter are not significantly different (post-ANOVA LSD; *P*=0.05).

Gene expression trends for up- and downregulated proteins followed a similar path in agroinfiltrated leaves compared to non-infiltrated leaves but could not explain the distinct accumulation patterns observed for some of these proteins in EV- and M2 vector-infiltrated leaves. For instance, mRNA transcript numbers for the two subunits of Rubisco decreased to similar levels in EV- and M2 vector-infiltrated leaves (**Fig. 7A**) but higher levels of both subunits were found in M2-expressing leaves (**Fig. 1, Fig. 6A**). Similarly, transcript numbers for the prominent PR-3 protein endochitinase A (Uniprot accession P08252) were comparable in EV- and M2 vector-infiltrated leaves (**Fig. 7B**) but PR-3 protein (including endochitinase A) levels were systematically lower upon M2 expression (**Fig. 6B**, **Supplemental Table S5**). A closer look at the proteome datasets in fact revealed a general trend for the accumulation of stress-related – including ER stress-associated and PR– proteins in agroinfiltrated leaves, by which the steady-state levels of these proteins in M2-expressing leaves, albeit greater than in non-infiltrated leaves, were only ∼20–70% the levels measured in EV-infiltrated leaves (**Table 1**). Overall, these data pointed to the onset of transcriptional and posttranscriptional regulatory events in M2-expressing leaves shaping, together, a defense-oriented proteome globally similar to, but nevertheless distinct from, the proteome of EV-infiltrated leaves.

**Figure 7.**
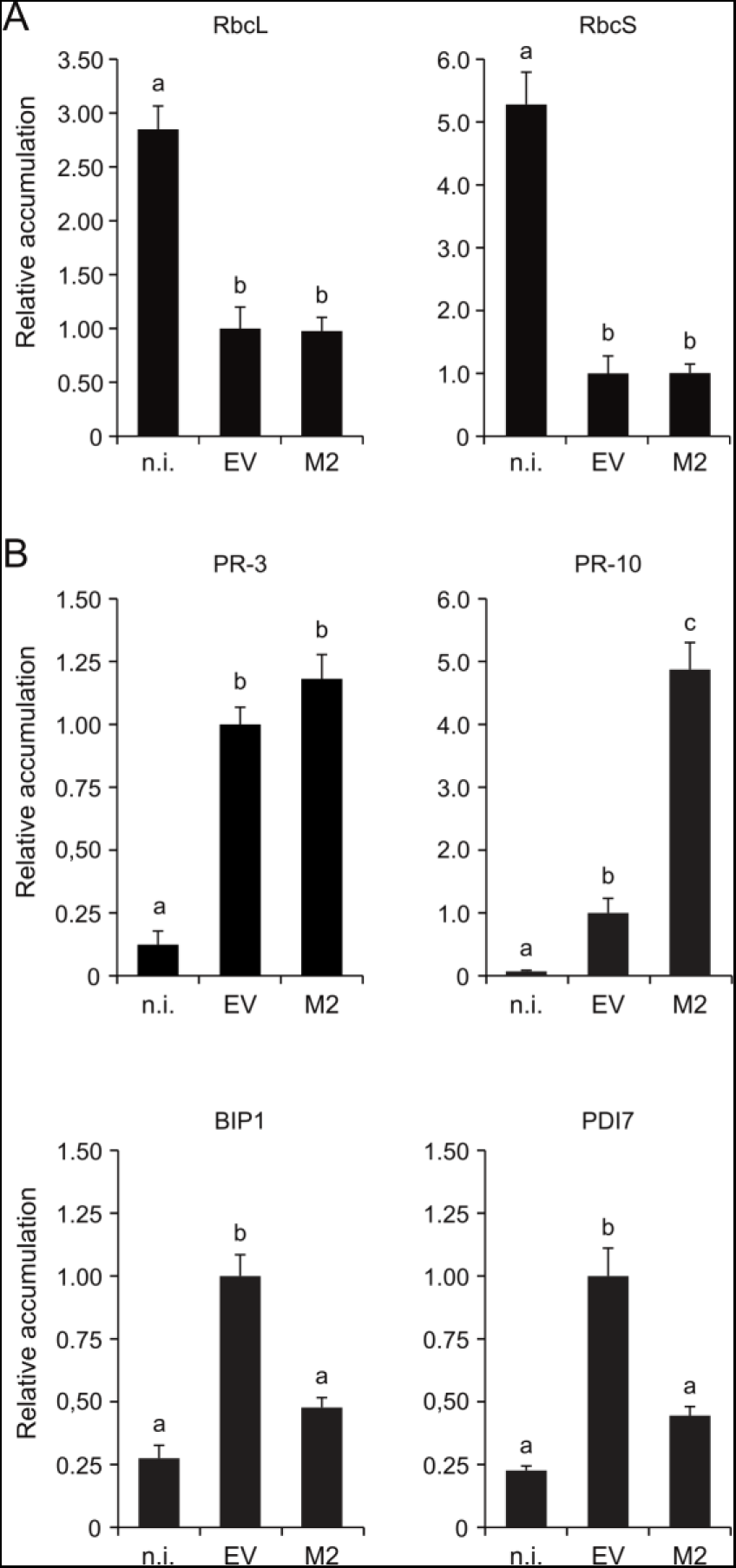
RT-qPCR analysis of Rubisco and defense-related protein transcripts in RNA extracts of non-infiltrated (n.i.), EV-infiltrated and M2-expressing leaves. (A) Relative abundance of transcripts for the large (RbcL) and small (RbcS) subunits of Rubisco. (B) Relative abundance of transcripts for PR-3 protein endochitinase A (UniProt Accession P08252), PR-10 protein (UniProt Accession A0A068JKR2), ER chaperone-associated protein BIP1 and protein disulfide isomerase PDI7. Data are expressed relative to EV-infiltrated leaves (arbitrary value of 1.0). Each bar is the mean of four biological (plant) replicate values ± se. Bars with the same letter are not significantly different (post-ANOVA LSD; *P*=0.05). Details on DNA primers for the qPCR amplifications are provided in **Supplemental Table S9**.

**Table 1.**
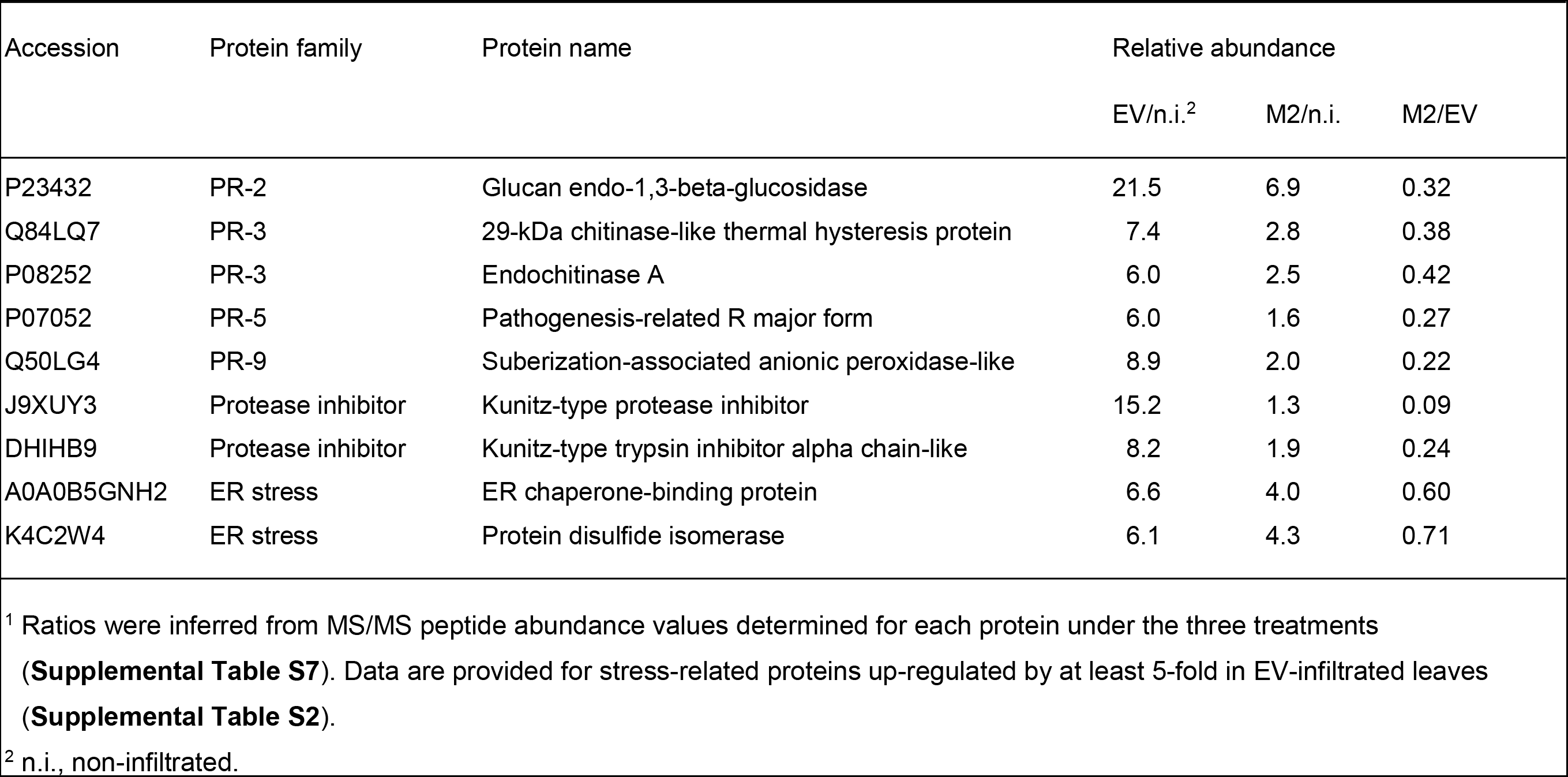
Relative abundance of stress-related proteins in EV-infiltrated [or M2-expressing] leaves compared to non-infiltrated [or EV-infiltrated] leaves^1^

| Accession | Protein family | Protein name | Relative abundance |  |  |
| --- | --- | --- | --- | --- | --- |
|  |  |  | EV/n.i.<sup>2</sup> | M2/n.i. | M2/EV |
| P23432 | PR-2 | Glucan endo-1,3-beta-glucosidase | 21.5 | 6.9 | 0.32 |
| Q84LQ7 | PR-3 | 29-kDa chitinase-like thermal hysteresis protein | 7.4 | 2.8 | 0.38 |
| P08252 | PR-3 | Endochitinase A | 6.0 | 2.5 | 0.42 |
| P07052 | PR-5 | Pathogenesis-related R major form | 6.0 | 1.6 | 0.27 |
| Q50LG4 | PR-9 | Suberization-associated anionic peroxidase-like | 8.9 | 2.0 | 0.22 |
| J9XUY3 | Protease inhibitor | Kunitz-type protease inhibitor | 15.2 | 1.3 | 0.09 |
| DHIHB9 | Protease inhibitor | Kunitz-type trypsin inhibitor alpha chain-like | 8.2 | 1.9 | 0.24 |
| A0A0B5GNH2 | ER stress | ER chaperone-binding protein | 6.6 | 4.0 | 0.60 |
| K4C2W4 | ER stress | Protein disulfide isomerase | 6.1 | 4.3 | 0.71 |

<sup>1</sup> Ratios were inferred from MS/MS peptide abundance values determined for each protein under the three treatments (**Supplemental Table S7**). Data are provided for stress-related proteins up-regulated by at least 5-fold in EV-infiltrated leaves (**Supplemental Table S2**).
<sup>2</sup> n.i., non-infiltrated.

### M2 Channel Expression Influences the Protein Secretion Profile of Agroinfiltrated Leaves

Basic reasons for the attenuation of defense (e.g. PR) protein levels and the establishment of a hybrid proteome in M2-expressing leaves remain to be understood. A first explanation could be related to the transcriptional downregulation of ER stress-associated proteins such as the chaperone-associated protein BIP1 or the protein disulfide isomerase PDI7 upon M2 channel expression (**Fig. 7B**), which in turn could have limited the efficiency of secreted protein folding and stability in transfected cells. A complementary explanation would be a general interfering effect of M2 channel activity on secreted protein trafficking and host plant defense responses. Several studies have documented the involvement of endomembrane protein trafficking pathways in plant immune responses (Inada et al., 2014; Wang et al., 2016), instrumental to ensure proper secretion of antimicrobial proteins in the apoplast and a rapid migration of pattern-recognition receptor (PRR) proteins towards the plasma membrane (Ben Khaled et al., 2015). The biological significance of intracellular protein trafficking upon microbial challenge is well illustrated on the plant side by the gene inducing effects of salicylic acid, that not only triggers the expression of defense (e.g. PR) proteins and PRR’s (Tateda et al., 2014) but also the expression of secretory pathway-associated proteins including ER-resident chaperones and co-chaperones, protein disulfide isomerases and the ER membrane receptor of signal [peptide] recognition particle (Wang et al., 2005). The importance of protein trafficking on the microbial side is illustrated by the production of protein effectors affecting biochemical functions of the host cell secretory pathway (Mukhtar et al., 2011; Weßling et al., 2014) and by the recently reported hijacking of host cell endocytic pathways by agrobacteria to facilitate the trafficking of their virulence factors (Li and Pan, 2017). Considering the importance of pH homeostasis for secreted proteins (Martinière et al., 2013; Jutras et al., 2018), alteration of the Golgi lumen pH by M2 could here have represented a disturbing factor in transfected cells affecting to some extent the processing, trafficking and/or secretion of PRR’s and PR proteins following agroinfiltration.

We characterized soluble protein profiles in the apoplast of non-infiltrated, EV-infiltrated and M2-expressing leaves to document the eventual impact of M2 on protein secretion (**Fig. 8**). Agroinfiltration was shown previously to trigger a strong upregulation of defense protein secretion in *N. benthamiana* leaves leading to a significant, 4-fold increase of soluble protein content in the apoplast (Goulet et al., 2010b). Accordingly, leaf agroinfiltration increased soluble protein content by 4- to 6-fold in the apoplast, from a baseline content of 0.10 mg protein.ml^−1^ in apoplast extracts of non-infiltrated leaves to mean contents of 0.40 to 0.56 mg.ml^−1^ in the extracts of agroinfiltrated leaves (anova; *P*=0.023). Increased apoplastic protein content upon agrobacterial challenge was associated with the secretion of ∼28-kDa and 34-kDa proteins (**Fig. 8A**) corresponding to the proteins of similar size immunodetected above with anti-PR-3 (endochitinase A) and anti-PR-2 protein antibodies (**Fig. 6B**). In line with iTRAQ and immunodetection data (**Fig. 3B, Fig. 6B**), the major band at 34 kDa was found in M2-expressing leaves at levels about half the corresponding levels in EV-infiltrated leaves (**Fig. 8B**) despite similar numbers of mRNA transcripts for endochitinase A in leaves under either treatment (**Fig. 7**). A post-transcriptional mitigating effect of M2 on protein release in the apoplast was further substantiated with GFP variant pHluorin as a recombinant protein model (**Fig. 8C**). We recently reported a positive effect of M2 on pHluorin accumulation in *N. benthamiana* leaves, by which the fluorescence emission rates and steady-state levels of this protein are increased by more than twofold in M2-expressing leaf tissue for a comparable level of pHluorin-encoding transcripts (Jutras et al., 2018). Total pHluorin content in leaf tissue–as inferred from fluorescence emission rates–was here also increased by more than twofold upon M2 expression, but pHluorin content in the apoplast was identical in leaves expressing this protein either alone or along with the viral channel. Together, these observations pointed to a posttranscriptional effect of M2 on host leaf cells altering to some extent the integrity, trafficking and/or secretion of endogenous (e.g. defense) and heterologous proteins during their migration towards the apoplast. A basic question from this point will be to find out where, in the secretory pathway, is M2 influencing the fate of secreted proteins. A practical question will be to determine the resulting output of these effects on the quality and yield of clinically-useful recombinant proteins targeted to the apoplast for proper processing and maturation.

**Figure 8.**
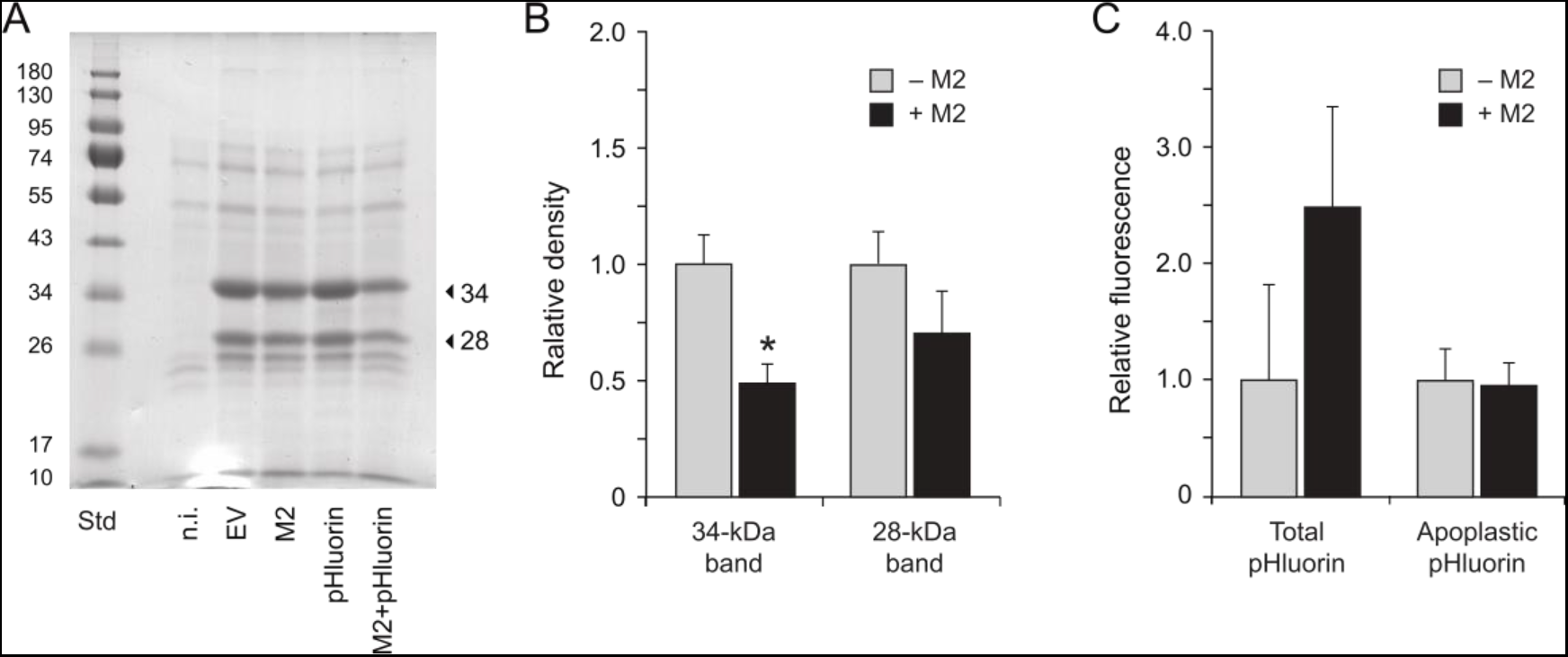
Protein secretion in the apoplast of non-infiltrated (n.i.), EV-infiltrated and M2 vector-infiltrated leaves expressing or not reporter protein pHluorin. (A) Apoplastic protein profiles following 12% w/v SDS-PAGE and Coomassie blue staining. Arrows point to the 28- and 34-kDa PR protein isoforms immunodetected above with anti-PR-2 and anti-PR-3 antibodies (**Fig. 6B**). Numbers on the left refer to commercial molecular weight standards (Std). (B) Abundance of the 28-kDa and 34-kDa PR proteins in apoplast protein preparations of agroinfiltrated leaves expressing (+M2) or not (−M2) the M2 channel. Abundance values were inferred from the volumic densities of Coomassie blue-stained 28-kDa and 34-kDa bands following SDS-PAGE (see panel A). Data are expressed relative to mean density values in apoplast extracts of ‘−M2’ (EV, pHluorin) infiltrated leaves. (C) Fluorescence emission at 520 nm in whole cell (Total) and apoplast protein extracts of agroinfiltrated leaves expressing pHluorin alone (−M2) or along with M2 (+M2). Data are expressed relative to fluorescence emitted in leaf extracts expressing pHluorin alone (−M2). Each bar is the mean of three biological (plant) replicates ± se.

## CONCLUSION

Our goal in this study was to look at the impact of influenza virus M2 proton channel expression on the proteome of agroinfiltrated *N. benthamiana* leaves, in an attempt to characterize the possible off-target effects of this accessory protein in a foreign protein production context. M2 channel expression was shown recently to trigger a partial neutralization of the Golgi lumen in *N. benthamiana* leaf cells helpful to stabilize pH-labile and protease-susceptible recombinant proteins in the cell secretory pathway (Jutras et al., 2015; 2018). We here followed a iTRAQ proteomics procedure to monitor proteome changes in M2-expressing leaves, keeping in mind the involvement of protein secretion in plant cells upon microbial challenge and the importance of pH homeostasis on protein maturation and trafficking in the secretory pathway. Our data pointed overall to a defense response-attenuating effect of M2 upon agroinfiltration, correlated with a restoration of Rubisco and soluble protein contents in leaf tissue. Studies will be welcome in coming years to assess the net impact of M2-induced proteome changes on recombinant protein yields *in planta.* The positive impact of M2 on the production of primary metabolism-associated proteins would suggest in practice an enhanced ability of the plant to accumulate recombinant proteins. The attenuation of defense protein secretion upon M2 expression could indicate by contrast an altered interaction between transfected leaf cells and the bacterial transgene vector compromising the ability of the host plant to efficiently process and secrete certain proteins of clinical interest. Work is underway to address these questions using recombinant protein models targeted to different cellular locations. Work is also underway to further assess the impact of M2 expression on secreted protein stability and integrity, given the influence of pH on endogenous protease activities in leaves (Jutras et al., 2018) and the possible retrofeedback effects of protease activity alterations on the leaf proteome (Badri et al., 2009; Goulet et al., 2010a).

## MATERIALS AND METHODS

### Transgene Constructs

Transgene constructs for pHluorin, M2 proton channel and M2 inactive mutant ^A30P^M2 were used as described previously (Jutras et al., 2015). Constructs harbored the appropriate DNA coding sequences fused to an upstream N-terminal signal peptide-encoding sequence for co-translational integration of the protein in the cell secretory pathway. The resulting coding sequences were assembled in a pCambia 2300 expression vector harboring an expression cassette for the silencing suppressor protein p19 (CAMBIA), between a duplicated Cauliflower mosaic virus 35S promoter in 5’ position and a nopaline synthase terminator sequence in 3’ position. An ‘empty’, pCambia 2300 vector was used as a positive control for the leaf agroinfiltrations. All vectors were maintained in *Agrobacterium tumefaciens*, strain AGL1 (Lazo et al., 1991) until use for the agroinfiltrations.

### Transient Expression in Leaves

Bacterial cultures for leaf infiltration were grown to stable phase in Luria-Bertani medium supplemented with appropriate antibiotics, and then harvested by gentle centrifugation at 4,000 *g* for 5 min at 20°C. The bacterial pellets were resuspended in 10 mM MES (2-[N-morpholino]ethanesulfonic acid), pH 5.6, containing 10 mM MgCl_2_ to an OD_600_ of 0.5, and incubated for 2 to 4 h at 20°C. Bacterial cultures harboring the M2-encoding vector, the ^A30P^M2-encoding vector or the empty vector were mixed at a volumic ratio of 1 in 4 with an EV- (or pHluorin vector)-harboring culture grown at the same optical density. The third leaves of 42 d-old plants (down from the main stem apex) were pressure-infiltrated with the bacterial suspensions using a needle-free syringe (D’Aoust et al., 2009), prior to plant incubation at 20°C in a Conviron PWG36 growth chamber (Conviron) for heterologous protein expression. Non-infiltrated plants were grown in parallel under the same conditions and used as negative controls. Leaf tissue was harvested 6 d post-infiltration for protein extraction and analysis. Three independent replicates each including the third leaf of three plants were used for each treatment to minimize variation of protein expression levels and to allow for statistical analysis of the data (Robert et al., 2013).

### Protein Extraction

Leaf tissue for whole-cell protein extraction was harvested as leaf discs representing 160 mg of infiltrated tissue and homogenized by disruption with ceramic beads (BioSpec) in a Mini-Bead beater apparatus (OMNI International). Total soluble proteins were extracted in three volumes of phosphate-buffered saline (PBS), pH 7.3, containing 5 mM EDTA, 0.5% w/v sodium deoxycholate (DOC) and 1 mM phenylmethylsulfonyl fluoride (Sigma-Aldrich). Leaf lysates were clarified by centrifugation for 20 min at 20,000 *g*, and total soluble proteins assayed according to Bradford (1976) with bovine serum albumin as a protein standard (Sigma-Aldrich). The resulting extracts were used directly for SDS-PAGE and immunoblotting, or stored at −20°C to reduce Rubisco levels before proteomic analysis (Qiu et al., 2008; Sainsbury et al., 2016). Leaf apoplast proteins were recovered as described (Robert et al., 2013), with some modifications. Freshly harvested leaves were washed in double-distilled water and submerged in agroinfiltration buffer (10 mM MES buffer, pH 5.8). Washed leaves were vacuum-infiltrated for 60 s at −80 kPa with infiltration buffer, dried off to remove excess buffer, rolled in a homemade Swiss-roll cylinder, and centrifuged at 4°C for 10 min at 1,000 *g* to collect the vacuum infiltrate. The resulting protein preparations were centrifuged at 6,000 *g* for 5 min at 4°C to discard *A. tumefaciens* cells. Protein content was assayed according to Bradford (1976) with bovine serum albumin as a protein standard, and the samples kept at −80°C until further use.

### iTRAQ Sample Preparation and Labeling

Whole-cell protein extracts (see above) from three biological replicates were used for iTRAQ proteomics. Similar volumes of the three replicates were pooled and the resulting mixture incubated overnight at −20°C in five volumes of pre-chilled acetone. Precipitated proteins were centrifuged at 20°C for 15 min at 16,000 g, and the protein pellets resuspended in 0.5 M triethylammonium bicarbonate (TEAB)−0.5% w/v sodium deoxycholate (DOC) following air drying at 20°C. Protein concentration in each sample was determined according to Bradford (1976), and 50 μg of protein was taken apart for iTRAQ labeling. TEAB and DOC were added to the samples at final concentrations of 0.5 M and 0.5% w/v, respectively. The proteins were reduced with Tris(2-carboxyethyl)phosphine (TCEP) and alkylated with methyl methanethiosulfonate (MMTS) according to the iTRAQ kit manufacturer’s instructions (Applied Biosystems), and then digested overnight at 37°C with sequence grade-trypsin (Promega) at a protease–protein ratio of 1:30. The resulting peptides were acidified to precipitate the DOC detergent, purified with an Oasis HLB cartridge (1 cc, 10 mg; Waters), SpeedVac-dried and dissolved in 30 μl of 0.5 M TEAB. Four-plex labeling was performed for 2 h in the dark at 20°C with the iTRAQ reagent (Applied Biosystems), and the labeled peptides combined in a single tube. The samples were SpeedVac-dried, cleaned up using an HLB cartridge (Waters) and separated in 14 fractions on a high pH (pH 10) reversed-phase chromatography column using the Agilent 1200 HPLC system (Agilent). Peptide fractions were SpeedVac-dried and resuspended in 0.1% v/v formic acid prior to MS/MS analysis.

### Mass Spectrometry

Peptide fractions containing approximately 900 ng of peptides were separated by online reversed-phase nanoscale capillary LC and analyzed by electrospray mass spectrometry. Separations were performed using a Dionex UltiMate 3000 nanoRSLC chromatography system (Thermo Fisher Scientific/Dionex Softron GmbH) connected to an Orbitrap Fusion mass spectrometer (Thermo Scientific) equipped with a nanoelectrospray ion source. The peptides were trapped in loading solvent (2% v/v acetonitrile, 0.05% v/v trifluoroacetic acid) for 5 min at 20 μL.min^−1^ on a 5 mm × 300 μm C18 pepmap cartridge pre-column (Thermo Fisher Scientific/Dionex Softron GmbH). The pre-column was switched online to a self-made 50 cm × 75 μm internal diameter separation column packed with ReproSil-Pur C18-AQ 3-μm resin (Dr. Maisch HPLC GmbH). The peptides were eluted over 90 min at 300 nL.min^−1^ along a 5–40% linear gradient of solvent B (80% v/v acetonitrile, 0.1% v/v formic acid) against 0,1% v/v formic acid (solvent A). Mass spectra were acquired under a data-dependent acquisition mode using the Thermo XCalibur software, v. 3.0.63. Full scan mass spectra in the 350–1800 *m/z* range were acquired in the Orbitrap spectrometer using an AGC target of 4e5, a maximum injection time of 50 ms, a resolution of 120,000 and an internal lock mass calibration on *m/z* 445.12003 (siloxane ion). Each MS scan was followed by acquisition of fragmentation MS/MS spectra of the most intense ions for a total cycle time of 3 s (top speed mode). Selected ions were isolated using a quadrupole analyzer in a window of 1.6 *m/z*, and fragmented by higher energy collision-induced dissociation with collision energy set at 45. Resulting fragments were detected in the Orbitrap at a resolution of 60,000, with an AGC target of 1e5 and a maximum injection time of 120 ms. Dynamic exclusion of previously fragmented peptides was set at a tolerance of 10 ppm for a period of 20 s.

### Database Searching

MS/MS spectra were analyzed with the Proteome Discover program, v. 2.1 (Thermo Scientific) set up to search the Solanaceae protein database of Uniprot (http://www.uniprot.org/taxonomy/4070; 117,836 proteins). Search parameters for matching were as follows: MMTS-alkylated Cys residues and iTRAQ-modified Lys, Tyr and peptide N-terminus as static modifications; oxidized Met residues and deamidated Asn and Gln residues as variable modifications; a mass search tolerance of 10 ppm for MS or 25 atomic mass units for MS/MS; and a maximum of two missed trypsin cleavages allowed. Protein identifications were deemed as valid when a False Discovery Rate of 1% was determined at the peptide and protein levels based on the target-decoy approach (Elias and Gygi, 2007). Protein Discoverer outputs were exported to the Microsoft Excel spreadsheet software, v. 2016 (Microsoft Inc.) for further analysis. A protein was considered as underexpressed when a ratio value of 0.5, or lower than 0.5, was calculated compared to the control; or as overexpressed when this ratio was equal to, or higher than, 2.0.

### BLAST Searches and GO Enrichment Analyses

The BLAST (https://blast.ncbi.nlm.nih.gov/Blast.cgi) and Blast2GO (https://www.blast2go.com) (Conesa et al., 2005) programs were used online to identify and classify the most downregulated, and most upregulated, proteins in leaves under the different experimental treatments. GO enrichment analyses were undertaken to compare the tested proteomes, based on the Gene Ontology system for gene and gene product classification (The Gene Ontology Consortium, 2008). A minimal E-value of 1 was set in Blast2GO for the BLASTP analysis, and the first 20 BLAST hits were selected for further analysis. A number of genes with no annotations in the custom database were annotated, wherever possible, using the GenBank UniProt database (http://www.uniprot.org). Predicted subcellular localization of the identified proteins was inferred using the Plant mPloc web server (http://www.csbio.sjtu.edu.cn/bioinf/plant-multi) (Chou and Shen, 2010). All DNA sequences were BLAST-searched against the *N. benthamiana* chloroplast genome (http://sefapps02.qut.edu.au) to identify chloroplastic proteins.

### Principal Component Analysis

A PCA was performed for the proteins confidently identified by MS/MS in protein extracts of non-infiltrated, EV-infiltrated and M2 channel-expressing leaves (i.e. 1,255 proteins overall). The relative abundance of each protein in each group was inferred from the iTRAQ MS/MS dataset (**Supplemental Table S1**). The PCA was performed on log-normalized data using the R software, v. 1.1.423 (R-Studio, www.rstudio.com). Graphical visualization of the PCA data was generated with the *ggbiplot* package.

### Immunoblotting and Protein Quantitation by Densitometry

Rubisco large (RbcL) and small (RbcS) subunits, PR-2 proteins and PR-3 proteins were detected by immunoblotting on nitrocellulose sheets following 12% (w/v) SDS-PAGE in reducing conditions. Rubisco subunits were detected with polyclonal IgG raised in rabbits against RbcL (Agrisera, Prod. No. AS03 037) or RbcS (AS07 259A). The PR proteins were detected with rabbit polyclonal IgG directed against PR-2 proteins (Agrisera, Prod. No. AS12 2366) or PR-3 proteins (no. AS07 207). Nonspecific binding on nitrocellulose sheets was prevented by incubation in blocking solution (5% w/v skim milk powder in PBS, containing 0.025% v/v Tween-20), which also served as antibody dilution buffer. The primary antibodies were detected with goat anti-rabbit secondary antibodies conjugated to alkaline phosphatase (Sigma-Aldrich). Protein signals were developed with the alkaline phosphatase substrate 5-bromo-4-chloro-3-indolyl phosphate and nitro blue tetrazolium as a colour indicator (Life Technologies). Densitometric analysis was performed using the Phoretix 2D Expression software v. 2005 (NonLinear USA) on non-saturated immunoblot images digitalized with an Amersham Image Scanner (GE Healthcare). All measurements were made with leaf extracts from at least three independent (plant) replicates.

### RNA Extraction and Quantification of mRNA Transcripts

RT-qPCR assays were performed with leaf samples from four plant replicates each harvested as two leaf discs representing 100 mg of fresh tissue. Leaf discs were ground in liquid nitrogen and total RNA extracted using the EZ-10 Spin Column Plant RNA Miniprep Kit (Biobasics). Residual DNA was removed using the RNase-free DNase Set (Qiagen) and RNA integrity assessed using an Agilent 2100 Bioanalyzer (Agilent Technologies). RNA quality was confirmed and concentration determined using a NanoDrop ND-1000 spectrophotometer (NanoDrop Technologies, Wilmington DE, U.S.A.), before reverse transcription to cDNA using the QuantiTect Reverse Transcription kit (Qiagen). Transcript quantification was performed by real-time RT-qPCR in 96-well plates using the ABI PRISM 7500 Fast real-time PCR system and custom data analysis software, version 2.0.1 (Thermo Fisher Scientific). Each PCR reaction contained 5 ng of cDNA template, 0.5 μM forward and reverse primers for target gene amplification (**Supplemental Table 9**) and 1X SYBR Green Master Mix (QuantiTect SYBR Green mix, Qiagen), for a total volume of 10 μL. qPCR was run under the SYBR Green amplification mode at the following PCR cycling conditions: 15 min incubation at 95°C, followed by 40 amplification cycles at 95°C for 5 s, 60°C for 30 s, and 65°C for 90 s. Reactions in the absence of cDNA template were conducted as negative controls and fluorescence readings were taken at the end of each cycle. The absence of DNA primer dimers and specificity of the amplifications were confirmed by melting curve analysis at the end of each reaction. Fluorescence and cycle threshold (Ct) values were exported to and analyzed using the Microsoft Excel spreadsheet software, v. 2016 (Microsoft, Inc.). The relative number of transcripts (1/2^Ct^) was averaged from technical RT-qPCR duplicates and used for subsequent normalization. Expression data were normalized against the geometric mean of six reference genes (**Supplemental Table S9**) to correct for biological variability and technical variations during RNA extraction, quantification and reverse transcription. Stability of reference gene expression was evaluated using the geNORM VBA applet for Microsoft Excel (Vandesompele et al., 2002). Fold changes in gene expression, reported relative to EV-infiltrated leaf tissue, were calculated using the 2^−ΔΔCt^ method (Livak and Schmittgen, 2001; Bustin et al., 2009). Standard deviation (sd) related to within-treatment biological variation was calculated in accordance with the error propagation rules.

### Recombinant pHluorin Quantification

pHluorin expression was monitored by detection of fluorescence emission with a Fluostar Galaxy microplate reader (BMG, Offenburg, Germany) using excitation and emission filters of 485 and 520 nm, respectively (Jutras et al., 2018). Fluorescence levels were expressed relative to fluorescence emission in M2-free (− M2) ‘control’ extracts. Samples were loaded in triplicate on Costar 96-well black polystyrene plates (Cedarlane, Burlington ON, Canada). All measurements were made with leaf protein extracts from six independent (plant) replicates.

### Statistical Analyses

Statistical analyses were performed using RStudio, v. 0.98.1103 (RStudio, Inc.). Analysis of variance (ANOVA) tests were used to compare peptide (protein) counts and mRNA transcript numbers among treatments. Contrast calculations and LSD mean comparison tests were performed for those ANOVA giving significant *P* values at an alpha value threshold of 5%.

## Supplemental Data

**Fig. S1** Complement to Fig. 1: Soluble protein content over 12 d in non-infiltrated leaves or agroinfiltrated leaves expressing or not the M2 proton channel

**Fig. S2** Complement to Fig. 2: GO enrichment analysis of iTRAQ-quantified proteins up- or downregulated in EV-infiltrated leaves compared to non-infiltrated control leaves

**Fig. S3** Complement to Fig. 3: GO enrichment analysis of iTRAQ-quantified proteins up- or downregulated in M2 vector-infiltrated leaves compared to EV-infiltrated leaves

**Table S1** Complete list of confidently identified proteins following iTRAQ analysis

**Table S2** The 60 most upregulated proteins in EV-infiltrated leaves compared to non-infiltrated control leaves

**Table S3** The 60 most downregulated proteins in EV-infiltrated leaves compared to non-infiltrated control leaves

**Table S4** The 60 most upregulated proteins in M2 vector-infiltrated leaves compared to EV-infiltrated leaves

**Table S5** The 60 most downregulated proteins in M2 vector-infiltrated leaves compared to EV-infiltrated leaves

**Table S6** The 100 most abundant proteins in non-infiltrated leaves

**Table S7** The 100 most abundant proteins in EV-infiltrated leaves

**Table S8** The 100 most abundant proteins in M2-channel expressing leaves

**Table S9** DNA primers for RT-qPCR amplifications

## ACKNOWLEDGMENTS

This work was supported by a Discovery grant from the Natural Science and Engineering Research Council (NSERC) of Canada to D.M. P.V.J. was the recipient of an AgroPhytoSciences NSERC–CREATE scholarship and of a BMP graduate scholarship co-funded by Medicago inc., NSERC and Québec Government’s research funding body FRQNT.

## LITERATURE CITED

Anand A, Uppalapati SR, Ryu CM, Allen SN, Kang L, Tang Y, Musore KS (2008) Salicylic acid and systemic acquired resistance play a role in attenuating Crown gall disease caused by *Agrobacterium tumefaciens*. Plant Physiol 146: 703–715

Ashnest JR, Huynh DL, Dragwidge JM, Ford BA, Gendall AR (2015) Arabidopsis intracellular NHX-type sodium-proton antiporters are required for seed storage protein processing. Plant Cell Physiol 56: 2220–2233

Badri MA, Rivard D, Coenen K, Michaud D (2009) Unintended molecular interactions in transgenic plants expressing clinically-useful proteins–The case of bovine aprotinin travelling the potato leaf cell secretory pathway. Proteomics 9: 746–756

Bassil E, Blumwald E (2014) The ins and outs of intracellular ion homeostasis: NHX-type cation/H^+^ transporters. Curr Opin Plant Biol 22: 1–6

Ben Khaled S, Robatzek S (2015) A moving view: subcellular trafficking processes in pattern recognition receptor-triggered plant immunity. Annu Rev Plant Biol 53: 379–402

Boller T, Felix G (2009) A renaissance of elicitors: perception of microbe-associated molecular patterns and danger signals by pattern recognition receptors. Annu Rev Plant Biol 60: 379–406

Bradford MM (1976) A rapid and sensitive method for the quantitation of microgram quantities of protein utilizing the principle of protein-dye binding. Anal Biochem 72: 248–254

Brewis IA, Brennan P (2010) Proteomics technologies for the global identification and quantification of proteins. Adv Prot Chem Struct Biol 80: 1–43

Bustin SA, Benes V, Garson JA, Hellemans J, Huggett J, Kubista M, Mueller R, Nolan T, Pfaffl MW, Shipley GL, et al. (2009) The MIQE guidelines: minimum information for publication of quantitative realtime PCR experiments. Clin Chem 55: 611–622

Cady SD, Luo W, Hu F, Hong M (2009) Structure and function of the Influenza A M2 proton channel. Biochemistry 48: 7356–7364

Chou KC, Shen HB (2010) Plant-mPLoc: A top-down strategy to augment the power for predicting plant protein subcellular localization. PLoS One 5: e11335

Conesa A, Götz S, García-Gómez JM, Terol J, Talón M, Robles M (2005) Blast2GO: a universal tool for annotation, visualization and analysis in functional genomics research. Bioinformatics 21: 3674–3676

D’Aoust MA, Lavoie PO, Belles-Iles J, Bechtold N, Martel M, Vézina LP (2009) Transient expression of antibodies in plants using syringe agroinfiltration. Meth Mol Biol 483: 41–50

Dettmer J, Hong-Hermesdorf A, Stierhof YD, Schumacher K (2006) Vacuolar H^+^-ATPase activity is required for endocytic and secretory trafficking in *Arabidopsis*. Plant Cell 18: 715–730

Elias JE, Gygi SP (2007) Target-decoy search strategy for increased confidence in large-scale protein identifications by mass spectrometry. Nat Meth 4: 207–214

Faye L, Boulaflous A, Benchabane M, Gomord V, Michaud D (2005) Protein modifications in the plant secretory pathway: current status and practical implications in molecular pharming. Vaccine 23: 1770–1778

Gomord V, Fitchette AC, Menu-Bouaouiche L, Saint-Jore-Dupas C, Plasson C, Michaud D, Faye L (2010) Plant-specific glycosylation patterns in the context of therapeutic protein production. Plant Biotechnol J 8: 564–587

Goulet C, Khalf M, Sainsbury F, D’Aoust MA, Michaud D (2012) A protease activity-depleted environment for heterologous proteins migrating towards the leaf cell apoplast. Plant Biotechnol J 10: 83–94

Goulet C, Benchabane M, Anguenot R, Brunelle F, Khalf M, Michaud D (2010a) A companion protease inhibitor for the protection of cytosol-targeted recombinant proteins in plants. Plant Biotechnol J 8: 142–154

Goulet C, Goulet C, Goulet MC, Michaud D (2010b) 2-DE proteome maps for the leaf apoplast of *Nicotiana benthamiana*. Proteomics 10: 2536–2544

Henkel JR, Weisz OA (1998) Influenza virus M2 protein shlows traffic along the secretory pathway. pH perturbation of acidified compartments affects early Golgi transport steps. J Biol Chem 273: 6518–6524

Holsinger LJ, Nichani D, Pinto LH, Lamb RA (1994) Influenza A virus M2 ion channel protein: a structure-function analysis. J Virol 68: 1551–1563

Ishiga Y, Watanabe M, Ishiga T, Tohge T, Matsuura T, Ikeda Y, Hoefgen R, Fernie AR, Mysore KS (2017) The SAL-PAP chloroplast retrograde pathway contributes to plant immunity by regulating glusosinolate pathway and phytohormone signaling. Mol Plant–Microbe Interact 30: 829–841

Inada N, Ueda T (2014) Membrane trafficking pathways and their roles in plant–microbe interactions. Plant Cell Physiol 55: 672–686

Jutras PV, Goulet MC, Lavoie PO, D’Aoust MA, Sainsbury F, Michaud D (2018) Recombinant protein susceptibility to proteolysis in the plant cell secretory pathway is pH-dependent. Plant Biotechnol J (in press): doi.org/10.1111/pbi.12928.

Jutras PV, Marusic C, Lonoce C, Deflers C, Goulet MC, Benvenuto E, Michaud D, Donini M (2016) An accessory protease inhibitor to increase the yield and quality of a tumour-targeting mAb in *Nicotiana benthamiana* leaves. PLoS One 11, e0167086

Jutras PV, D’Aoust MAA, Couture MMJ, Vézina LP, Goulet MC, Michaud D, Sainsbury F (2015) Modulating secretory pathway pH by proton channel co-expression can increase recombinant protein stability in plants. Biotechnol J 10: 1478–1486

Lazo GR, Stein PA, Ludwig RA (1991) A DNA transformation-competent Arabidopsis genomic library in *Agrobacterium*. Biotechnology 9: 963–967

Li X, Pan SQ (2017) Agrobacterium delivers VirE2 protein into host cells via clathrin-mediated endocytosis. Sci Adv 3: e1601528

Livak KJ, Schmittgen TD (2001) Analysis of relative gene expression data using real-time quantitative PCR and the 2^(−ΔΔC(T))^ method. Methods 25: 402–408

Lomonossoff GP, D’Aoust MA (2016) Plant-produced biopharmaceuticals: A case of technical developments driving clinical deployment. Science 353: 1237–1240

Makhzoum A, Benyammi R, Koustafa, K, Trémouillaux-Guiller J (2014) Recent advances on host plants and expression cassettes’ structure and function in plant molecular pharming. BioDrugs 28: 145–159

Mandal MK, Ahvari H, Schillberg S, Schiermeyer A (2016) Tackling unwanted proteolysis in plant production hosts used for molecular farming. Front Plant Sci 7: 1–6

Martinière A, Bassil E, Jublanc E, Alcon C, Reguera M, Sentenac H, Blumwald E, Paris D (2013) In vivo intracellular pH measurements in tobacco and *Arabidopsis* reveal an unexpected pH gradient in the endomembrane system. Plant Cell 25: 4028–4043

Mukhtar MS, Carvunis AR, Dreze M, Epple P, Steinbrenner J, Moore J, Tasan M, Galli M, Hao T, Nishimura MT, Pevzner SJ, Donovan SE, Ghamsari L, Santhanam B, Romero V, Poulin MM, Gebreab F, Gutierrez BJ, Tam S, Monachello D, Boxem M, Harbort CJ, McDonald N, Gai L, Chen H, He Y, European Union Effectoromics Consortium, Vandenhaute J, Roth FP, Hill DE, Ecker JR, Vidal M, Beynon J, Braun P, Dangl JL (2011) Independently evolved virulence effectors converge onto hubs in a plant immune system network. Science 3111: 596–601

Nagatoshi Y, Ikeda M, Kishi H, Hiratsu K, Muraguchi A, Ohme-Takagi M (2015) Induction of a dwarf phenotype with IBH1 may enable increased production of plant-made pharmaceuticals in plant factory conditions. Plant Biotechnol J 14: 887–894

Nomura H, Komori T, Uemura S, Kanda Y, Shimotani K, Nakai K, Furuichi T, Takebayashi K, Sugimoto T, Sano S, Suwastika IN, Fukusaki E, Yoshioka H, Nakahira Y, Shiina T (2012) Chloroplast-mediated activation of plant immune signalling in *Arabidopsis*. Nat Commun 3: 926

Orlowski J, Grinstein S (2011) Na+/H+ exchangers. Comprehensive Physiol 1: 2083–2100

Pillay P, Schlüter U, van Wyk S, Kunert KJ, Vorster BJ (2014) Proteolysis of recombinant proteins in bioengineered plant cells. Bioengineered 5: 1–6

Pruss GJ, Nester EW, Vance V (2008) Infiltration with *Agrobacterium tumefaciens* induces host defense and development-dependent responses in the infiltrated zone. Mol Plant-Microbe Interact 21: 1528–1538

Qiu QS, Huber JL, Booker FL, Jain V, Leakey ADB, Fiscus EL, Yau PM, Ort DR, Huber SC (2008) Increased protein carbonylation in leaves of *Arabidopsis* and soybean in response to elevated [CO_2_]. Photosynth Res 97: 155–166

Reguera M, Bassil E, Tajima H, Wimmer M, Chanoca A, Otegui MS, Paris N, Blumwald E (2015) pH regulation by NHX-type antiporters is required for receptor-mediated protein trafficking to the vacuole in Arabidopsis. Plant Cell 27: 1200–1217

Robert S, Jutras PV, Khalf M, D’Aoust MA, Goulet MC, Sainsbury F, Michaud D (2016) Companion protease inhibitors for the *in situ* protection of recombinant proteins in plants. Meth Mol Biol 1385: 115–126

Robert S, Goulet MC, D’Aoust MA, Sainsbury F, Michaud D (2015) Leaf proteome rebalancing in *Nicotiana benthamiana* for upstream enrichment of a transiently expressed recombinant protein. Plant Biotechnol J 13: 1169–1179

Robert S, Khalf M, Goulet MC, D’Aoust MA, Sainsbury F, Michaud D (2013) Protection of recombinant mammalian antibodies from development-dependent proteolysis in leaves of *Nicotiana benthamiana*. PLoS One 8: e70203

Sainsbury F, Jutras PV, Vorster J, Goulet MC, Michaud D (2016) A chimeric affinity tag for efficient expression and chromatographic purification of heterologous proteins from plants. Front Plant Sci 7: 141

Sack M, Hofbauer A, Fischer R, Stoger E (2015) The increasing value of plant-made proteins. Curr Opin Biotechnol 32: 163–170

Sakaguchi T, Leser GP, Lamb RA (1996) The ion channel activity of the Influenza virus M_2_ protein affects transport through the Golgi apparatus. J Cell Biol 133: 733–747

Schnell JR, Chou JJ (2008) Structure and mechanism of the M2 proton channel of influenza A virus. Nature 451: 591–595

Schumacher K (2014) pH in the plant endomembrane system–an import and export business. Curr Opin Plant Biol 22: 71–76

Serrano I, Audran C, Rivas S (2016) Chloroplasts at work during plant innate immunity. J Exp Bot 67: 3845–3854

Shen J, Zeng Y, Zhuang X, Sun L, Yao X, Pimpl P, Jiang L (2013) Organelle pH in the *Arabidopsis* endomembrane system. Mol Plant 6: 1419–1437

Shimizu T, Satoh K, Kikuchi S, Omura T (2007) The repression of cell wall- and plastid-related genes and the induction of defense-related genes in rice plants infected with *Rice dwarf virus*. Mol Plant–Microbe Interact 3: 247–254

Stöger E, Fischer R, Moloney M, Ma JKC (2014) Plant molecular pharming for the treatment of chronic and infectious diseases. Annu Rev Plant Biol 65: 743–768

Streatfield SJ (2007) Approaches to achieve high-level heterologous protein production in plants. Plant Biotechnol J 5: 2–16

Sugano S, Jiang CJ, Miyazawa SI, Masumoto C, Yazawa K, Hayashi N, Shimono M, Nakayama A, Miyao M, Takatsuji H (2010) Role of OsNPR1 in rice defense program as revealed by genome-wide expression analysis. Plant Mol Biol 74: 549–562

Takatsuji H (2017) Regulating tradeoffs to improve rice production. Front Plant Sci 8: 171

Tateda C, Zhang Z, Shrestha J, Jelenska J, Chinchilla D, Greenberg JT (2014) Salicylic acid regulates *Arabidopsis* microbial pattern receptor kinase levels and signaling. Plant Cell 26: 4171–4187

The Gene Ontology Consortium (2008) The Gene Ontology project in 2008. Nucl Acids Res 36: D440–444

Tschofen M, Knopp D, Hood E, Stöger E (2016) Plant molecular farming: much more than medicines. Annu Rev Plant Biol 9: 271–294

Vandesompele J, De Preter K, Pattyn F, Poppe B, Van Roy N, De Paepe A, Speleman F (2002) Accurate normalization of real-time quantitative RT-PCR data by geometric averaging of multiple internal control genes. Genome Biol 3: 34

Wang D, Weaver ND, Kesarwani M, Dong X (2005) Induction of protein secretory pathway is required for systemic acquired resistance. Science 308: 1036–1040

Wang WM, Liu PQ, Xu YJ, Xiao S (2016) Protein trafficking during plant innate immunity. J Integr Plant Biol 58: 284–298

Weßling R, Epple P, Altmann S, He Y, Yang L, Henz SR, McDonald N, Wiley K, Bader KC, Gläßer C, Mukhtar MS, Haigis S, Ghamsari L, Stephens AE, Ecker JR, Vidal M, Jones JDG, Mayer KFX, Ver Loren van Themaat E, Weigel D, Schulze-Lefert P, Dangl JL, Panstruga R, Braun P (2014) Convergent Targeting of a Common Host Protein-Network by Pathogen Effectors from Three Kingdoms of Life. Cell Host Micr 16: 364–375

Wilbers RHP, Westerhof RB, van Raaij DR, van Adrichem M, Prakasa AD, Lozano-Torres JL, Bakker J, Smant G, Schots A (2016) Co-expression of the protease furin in Nicotiana benthamiana leads to efficient processing of latent transforming growth factor-β1 into a biologically active protein. Plant Biotechnol J 14: 1695–1704

Wu X, Ebine K, Ueda T, Qiu QS (2016) AtNHX5 and AtNHX6 are required for the subcellular localization of the SNARE complex that mediates the trafficking of seed storage proteins in *Arabidopsis*. PLoS One 11: e0151658

Xiao Y, Savchenko T, Bidoo EEK, Chehab WE, Hayden DM, Tolstikov V, Corwin JA, Kliebenstein DJ, Keasling JD, Dehesh K (2012) Retrograde signaling by the plastidial metabolite MEcPP regulates expression of nuclear stress-response genes. Cell 149: 1525–1535

